# Breathing-Driven Modulation of Reticulospinal Tract Activity

**DOI:** 10.1101/2025.03.02.641029

**Authors:** Ruqayya Thawer, Stuart N Baker, Boubker Zaaimi

## Abstract

The reticulospinal tract (RST) plays a pivotal role in motor control, especially during recovery after neurological injuries such as stroke and spinal cord injury (SCI). Understanding how RST activity is modulated offers valuable insights into improving motor function recovery. Recent studies have demonstrated that breathing rhythms influence brain activity. This study explores how respiratory rhythms modulate RST excitability during motor tasks, using the StartReact paradigm to examine reaction times (RTs) across visual (VRT), visual-auditory (VART), and visual-auditory startling (VSRT) conditions. We measured RTs in three muscles (first dorsal interosseous, flexor digitorum superficialis, and biceps) in healthy adult participants (n=13, both sexes) performing multi-joint movements. RTs were longest in the VRT condition and significantly decreased when auditory stimuli were added (VART), with further reductions observed in the VSRT condition. Additionally, respiratory phase transitions, particularly from inspiration to expiration (IE), significantly influenced RTs, with the shortest RTs observed during these transitions in the VSRT condition. These findings suggest that RST excitability is dynamically modulated by respiratory rhythms. This modulation of the RST by respiratory phase transitions could inform future neurorehabilitation strategies, such as respiratory-phase-aligned stimulation, to enhance motor recovery following corticospinal lesions. Ultimately, this approach may optimize the timing of interventions, improving outcomes in conditions such as stroke and SCI.

**Significance Statement:** Brainstem pathways play a crucial role in motor recovery after stroke, and understanding how these pathways change during recovery is key to optimizing their participation in rehabilitation. This study demonstrates how respiratory rhythms influence these brainstem pathways. Using the StartReact paradigm, we show that muscle response times are faster when transitioning from inspiration to expiration. These findings suggest that the body’s natural breathing rhythms can enhance motor output by activating these pathways. This could inform innovative rehabilitation strategies, such as aligning interventions with specific respiratory phases, to improve motor recovery in stroke and spinal cord injury. Our research highlights the potential for personalized therapies that harness the body’s intrinsic rhythms to optimize recovery.

## Introduction

Stroke is a leading cause of long-term disability, often resulting in severe motor impairments (Katan and Luft, 2018). While therapies aimed at motor recovery have traditionally focused on rehabilitative training (Dee et al., 2020), neuromodulation techniques such as vagus nerve stimulation (VNS) are gaining attention for enhancing motor recovery (Ting et al., 2021; Keser et al., 2023; Huang et al., 2024). The mechanism underlying VNS-induced motor recovery remain unclear (Bowles et al., 2022; Andalib et al., 2023; Keser et al., 2023). How can a nerve primarily involved in regulating autonomic functions such as breathing and the transition between inspiration and expiration contribute to stroke-related motor recovery? One possibility is that subcortical motor circuits, particularly the reticulospinal tract (RST), may mediate VNS-driven improvements through their role in motor control and plasticity.

The reticular formation and its descending motor pathway, the RST, play a crucial role in motor control (Davidson and Buford, 2006; Riddle et al., 2009; Ortiz-Rosario et al., 2014). Our prior research has demonstrated that RST markedly increases its contribution to spinal motoneurons, compensating for lost corticospinal input (Zaaimi et al., 2012). However, this RST input is imbalanced, disproportionately favoring flexor over extensor motor neurons. Such imbalanced plasticity contributes to characteristic post-stroke motor impairments, including extensor weakness and flexion synergies. Additionally, we have shown that the reticular formation undergoes significant reorganization after corticospinal tract injury, with altered neuronal firing (Zaaimi et al., 2018b). Recent studies have highlighted the potential of targeting the RST to improve motor function after stroke (Choudhury et al., 2020). However, the factors that influence RST activity and how they can be harnessed for therapeutic purposes remain poorly understood.

Respiration, a fundamental physiological process, is increasingly recognized as a modulator of brain function. It has been demonstrated that breathing drives oscillations of the membrane potential of neurons in various cortical areas including prefrontal and somatosensory cortices (Juventin et al., 2023). A study (Li and Rymer, 2011) examined the relationship between respiration and motor activity by using TMS or electrical stimulation on hand muscles and discovered a general enhancement of the motor system related to breathing. The authors observed that delivering stimulation to finger extensors during voluntary inspiration resulted in a significant decrease in finger flexor spasticity in a stroke patient.

The transition from inspiration to expiration and vice versa is critical because these shifts in the body’s physiological state can modulate cortical excitability. A study by Zelano et al. (Zelano et al., 2016) demonstrated that the act of inhaling through the nose significantly increases delta oscillations and enhances cognitive functions such as memory recall, directly linking respiratory phase to cortical activity. Conversely, Nakamura et al. (Nakamura et al., 2018) suggest that the transition from expiration to inspiration might affect reaction time and recall accuracy in a delayed match-to-sample visual recognition task. Their results showed that when this transition occurred between image presentation and response, subjects experienced significantly longer response times and decreased recall accuracy compared to trials where this transition did not occur.

To probe the relationship between respiration and RST function, we turn to the StartReact paradigm. The StartReact paradigm, which involves a shortened reaction time to an unexpected auditory stimulus (Valls-Solé et al., 1995; Valls-Solé et al., 1999), provides a valuable tool for investigating subcortical motor pathways (Carlsen et al., 2004; Carlsen et al., 2011; Carlsen et al., 2012; Baker and Perez, 2017). If the therapeutic effect of VNS stems from leveraging respiration-driven modulation of RST excitability, we hypothesize that the transition from inspiration to expiration—a critical phase regulated by the vagus nerve’s inspiratory off-switch function (von Euler and Trippenbach, 1975)—could represent a period of heightened RST output. Consequently, we expect shorter reaction times for startling auditory stimuli occurring during these respiratory transition phases.

## Methods

### Participants

Thirteen healthy participants (age range: 20–22; 3 males, 10 females) were recruited for this study. Eligibility was determined through a pre-screening questionnaire to exclude individuals with any history of neurological disorders. All participants provided written informed consent before the commencement of the study, which was approved by the Human Research Ethics Committee at Aston University.

### Electromyography (EMG)

Participants were seated comfortably in a soundproof room designed to meet clinical auditory standards. EMG signals were recorded from the right arm’s biceps using adhesive surface electrodes (model H59P Kendall; Covidien, Dublin, Ireland). Electrodes were positioned over the flexor digitorum superficialis (FDS), first dorsal interosseous (1DI), and biceps muscles. For the FDS and biceps, the electrodes were spaced 2–3 cm apart, while for the 1DI, the spacing was 1 cm. Signals were amplified using a D360 amplifier (Digitimer Ltd, Welwyn Garden City, UK) with a gain of 1000 and a bandpass filter ranging from 30 Hz to 2 kHz. The amplified signals were digitized via a Micro1401 interface and recorded using Spike2 software (Cambridge Electronic Design, Cambridge, UK).

### Breathing Measurements

Breathing signals were measured using a surgical mask equipped with a temperature sensor (LM34, Texas Instruments) positioned below the nostrils. The mask was adjusted to avoid direct skin contact while ensuring comfort for the participants. The sensor’s signals were amplified with a gain of 500 and bandpass filtered between 0.3 and 3.4 Hz using a custom-built circuit. The filtered signals were then digitized via the Micro1401 interface for further analysis.

### StartReact Protocol

The StartReact protocol was adapted from the methods described by Baker and Perez (Baker and Perez, 2017). A red LED positioned one meter away from the participants served as the visual cue. Participants rested their right forearm on an armrest at a 90-degree angle and were instructed to perform a power grip and flex their elbow and wrist as quickly as possible in response to the LED flash, which lasted for 20 ms. Three conditions were presented in randomized order: a visual reaction time (VRT) condition using the LED light only, a visual-auditory reaction time (VART) condition combining the LED light with a quiet sound (80 dB, 500 Hz, 20 ms), and a visual-startle reaction time (VSRT) condition combining the LED light with a startling sound (115 dB, 500 Hz, 20 ms). Reaction times were measured as the interval between the LED cue and the onset of EMG activity.

Participants were familiarized with the task before the main experiment and instructed on all conditions to ensure proper understanding. To acclimate them to the startling stimulus, five loud sounds were presented at 5-second intervals without requiring any task performance. The experiment consisted of four blocks, with each block containing 20 trials for each condition (VRT, VART, VSRT) in random order. The intervals between trials varied randomly between 12.5 and 15.7 seconds. Participants were provided a 5-minute break between blocks to minimize fatigue.

### Respiratory Phase Categorization

The respiratory signals recorded during the experiment were segmented into inspiration and expiration periods. Each period was further divided into three equidistant phases, resulting in a total of six phases: the start (I1), middle (I2), and end of inspiration (I3), as well as the start (E1), middle (E2), and end of expiration (E3). Stimuli delivered during the experiment were categorized based on the respiratory phase at the time of occurrence to examine phase-specific effects on reaction times. Figure 1 illustrates the experimental setup and the process of respiratory signal analysis.

**Figure 1:**
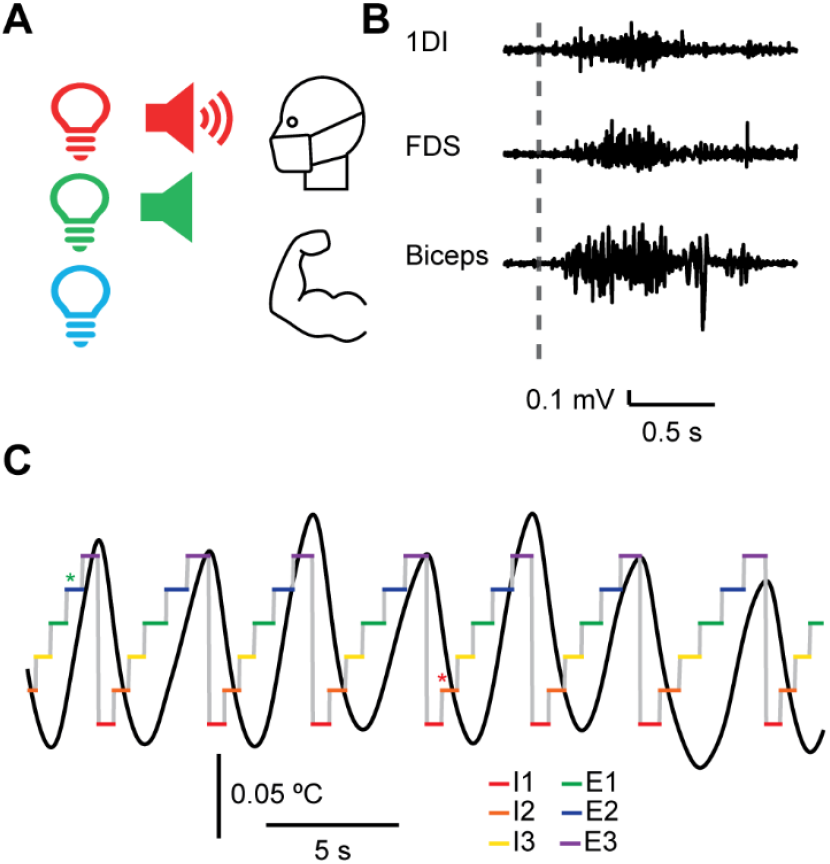
Recording of the StartReact experiment. **(A)** The participant is equipped with a mask containing a sensor to monitor respiratory signals and is instructed to perform an elbow and a power grip as soon as the LED light is activated. The light may be delivered alone (blue condition), accompanied by a quiet sound (green condition), or by a loud, startling sound (red condition). **(B)** All conditions are presented in a randomized order while recording the respiratory signal, as well as EMG activity from the 1DI, FDS, and Biceps muscles. **(C)** The respiratory signal (black trace) is segmented into six distinct phases, with a decrease in temperature during inspiration and an increase during expiration. Both the inspiration and expiration cycles are divided into three equidistant phases: start of inspiration (red, I1), middle of inspiration (orange, I2), end of inspiration (yellow, I3), start of expiration (green, E1), middle of expiration (blue, E2), and end of expiration (magenta, E3). Stimulations are categorized according to the respiratory phase in which they occur. In this example, a VART stimulation (green star) occurs during the middle of expiration (E2), while a VSRT stimulation (red star) occurs during the middle of inspiration (I2).

### Analysis

Reaction times for the three conditions (VRT, VART, VSRT) were assessed for normality using the Kolmogorov-Smirnov test in MATLAB. As the data were not normally distributed, non-parametric tests were employed. Differences across conditions were analyzed using the Mann-Whitney U test, while VART-VSRT differences from zero were assessed using the one-sample Wilcoxon signed-rank test.

To assess whether peaks and troughs in reaction times across respiratory phases were statistically significant, a model-based approach was used. We fitted a model made up of a mixture of two sinusoids to the reaction time data (*T*) as a function of the phase of respiration *x* (varying from 1 to 6; 1 indicates the onset of inspiration, and 4 the onset of expiration):

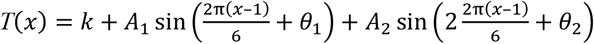

Here *k* is a constant offset, *A*_*1*_ and *A*_*2*_ the sinusoid amplitudes, and *θ*_*1*_ and *θ*_*2*_ are phase shifts (in radians).

Stimuli were grouped based on where they occurred within the respiratory phase, and phase-selective averages of reaction times measured, together with the standard error of the mean (SEM). The StartReact effect was calculated as the difference between the VART and VSRT for each respiratory phase and each muscle. To evaluate the significance of observed peaks and troughs in individual reaction time measures, or the StartReact effect, simulations were performed. A bootstrapping approach was employed to generate 10,000 simulated datasets by randomly shuffling reaction times across respiratory phases. For each simulated dataset, sinusoidal fits were computed. Peaks at the transition phases between inspiration and expiration (I3–E1) and troughs at the middle of inspiration (I2) and middle of expiration (E2) were extracted from the sinusoidal fits to both the real and simulated data. These phases were selected based on previous literature suggesting that motor excitability is particularly sensitive to changes during the transition from inspiration to expiration, where respiratory-related modulation is most pronounced. Conversely, a relative suppression of effects could be expected during the middle of inspiration or expiration, potentially reflecting more stable respiratory-driven motor states. The actual peak or trough was compared to the simulated distributions, and the p-value was calculated as the proportion of simulations that exceeded (for peaks) or fell below (for troughs) the actual value.

## Results

Figure 2A displays the average rectified EMG traces for the 1DI, FDS, and biceps muscles across the VRT, VART, and VSRT conditions for a single subject. In all muscles, the VSRT condition showed the shortest reaction times compared to both the VRT and VART conditions, indicating that the VSRT condition produced faster motor responses.

**Figure 2.**
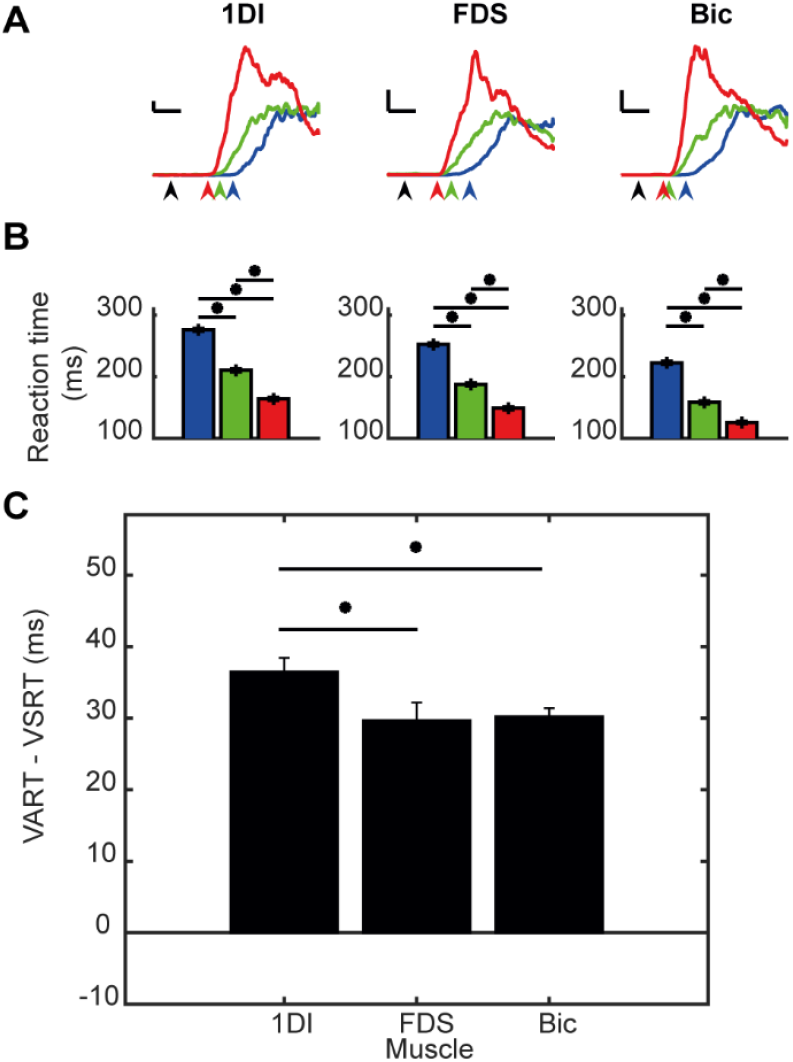
Rectified EMG traces, reaction times, amplitudes, and differences for VRT, VART, and VSRT conditions across muscles. **(A)** Rectified EMG traces for the 1DI (left), FDS (middle), and biceps (right) muscles during the VRT (blue), VART (green), and VSRT (red) conditions in a representative participant. Reaction times shortened during the VSRT condition compared with the VART and VRT conditions across all muscles. The black arrows are at 0 s (time of cue), and the blue, green, and red arrows show the reaction times for the VRT, VART, and VSRT, respectively. Calibration bars are: horizontal 100 ms, vertical 0.05 mV. **(B)** Bar charts showing reaction times (RT) for the VRT, VART, and VSRT conditions across the 1DI, FDS, and biceps muscles. Error bars represent the standard error of the mean (SEM). Asterisks indicate statistically significant differences based on the Mann-Whitney U test (*p < 0.05). **(C)** VART-VSRT differences across muscles, presented as bar plots with SEM error bars. A one-sample Wilcoxon signed-rank test confirmed that VART-VSRT differences were significantly greater than zero for all muscles. Pairwise comparisons between muscles were performed using the Mann-Whitney U test, with significant differences indicated by asterisks (*p < 0.05).

Figure 2B shows that for 1DI, reaction times were significantly (Mann Whitney U test) shorter in the VSRT condition compared to both VART (p < 0.01, 164.11 ± 66.44 ms vs. 210.37 ± 71.43 ms) and VRT (p < 0.01, 164.11 ± 66.44 ms vs. 276.04 ± 59.85 ms). Similarly, VART was significantly shorter than VRT (p < 0.01, 210.37 ± 71.43 ms vs. 276.04 ± 59.85 ms). For FDS, reaction times were shortest in the VSRT condition compared to VART (p < 0.01, 148.98 ± 65.44 ms vs. 187.51 ± 68.66 ms) and VRT (p < 0.01, 148.98 ± 65.44 ms vs. 252.39 ± 61.77 ms). VART was also shorter than VRT (p < 0.01, 187.51 ± 68.66 ms vs. 252.39 ± 61.77 ms). For the biceps, reaction times were again shortest in the VSRT condition compared to VART (p < 0.01, 125.69 ± 52.50 ms vs. 158.41 ± 64.60 ms) and VRT (p < 0.01, 125.69 ± 52.50 ms vs. 222.55 ± 65.04 ms). VART was also shorter than VRT (p < 0.01, 158.41 ± 64.60 ms vs. 222.55 ± 65.04 ms).

Figure 2C illustrates the results of the VART-VSRT difference analysis across muscles. The one-sample Wilcoxon signed-rank test confirmed that the VART-VSRT difference was significantly different from zero for all three muscles: 1DI (Mean = 36.43, SEM = 1.99, p = 0.0002), FDS (Mean = 29.61, SEM = 2.59, p < 0.01), and biceps (Mean = 30.16, SEM = 1.23, p < 0.01). Pairwise comparisons between muscles using the Mann-Whitney U test revealed significant differences between 1DI and FDS (p = 0.016) as well as between 1DI and biceps (p = 0.006), while the difference between FDS and biceps was not significant (p = 0.442). These findings suggest distinct VART-VSRT dynamics, with the 1DI showing the largest effect.

Figure 3 illustrates sinusoidal fits modelling the modulation of RTs across six respiratory phases for the three experimental conditions. The six respiratory phases represent the start, middle, and end of inspiration (I1, I2, and I3) and expiration (E1, E2, and E3), with the inspiration-expiration transition occurring between I3 and E1.

**Figure 3:**
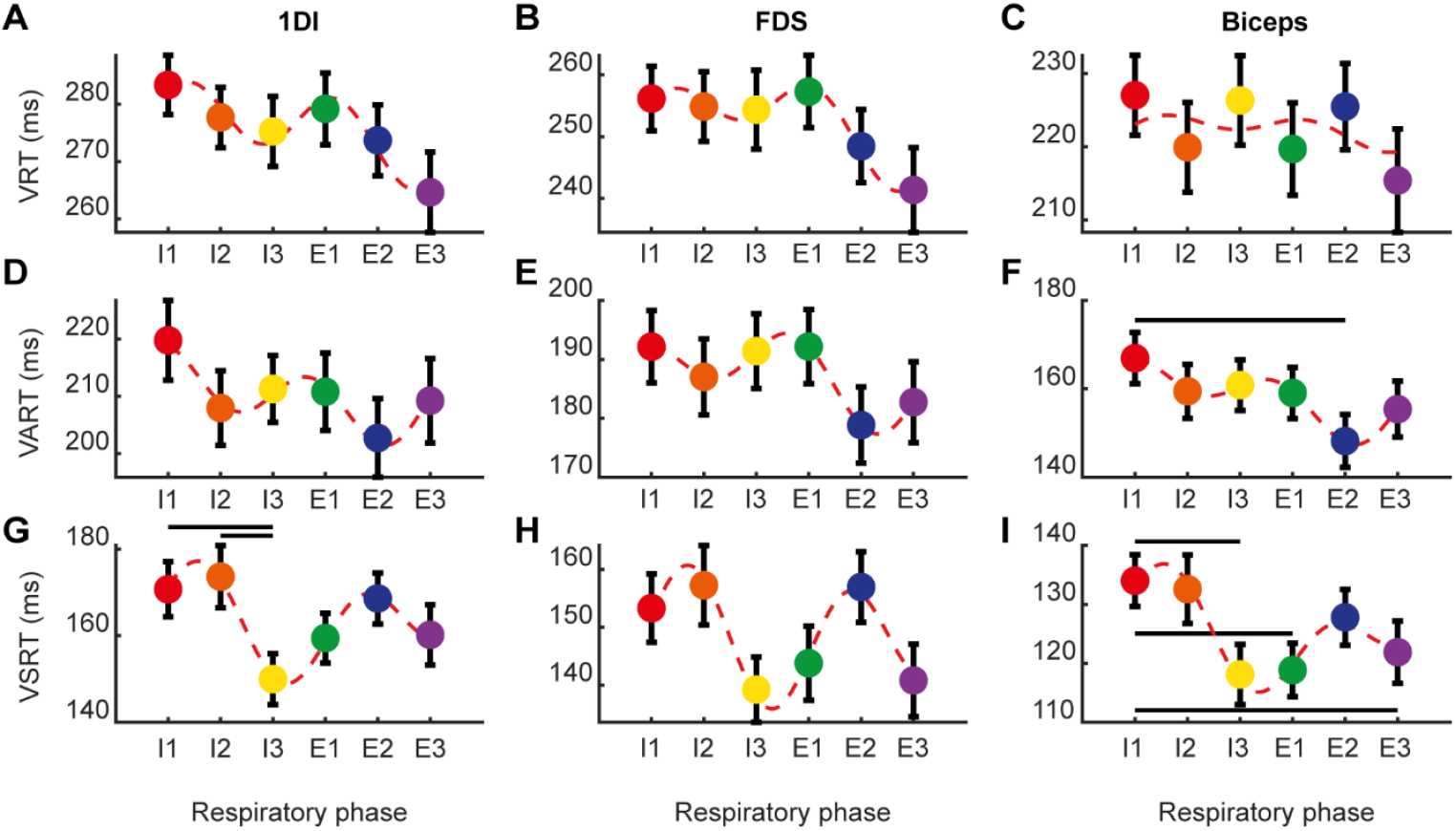
Sinusoidal fits illustrating modulation of reaction times across respiratory phases. The respiratory phases are defined with the inspiration phases (I1-3) and expiration phases (E1-3) as follows: I1 (red, start of inspiration), I2 (orange, middle of inspiration), I3 (yellow, end of inspiration), E1 (green, start of expiration), E2 (blue, middle of expiration), and E3 (magenta, end of expiration). **(A-C)** show the sinusoidal fits for the 1DI, FDS, and biceps muscles in the VRT condition, with minimal modulation and no significant differences in reaction times across the respiratory cycle. **(D-F)** illustrate the modulation in the VART condition, showing slight enhancements in reaction times across the respiratory phases, though these changes were not statistically significant. **(G-I)** depict the strongest modulation in the VSRT condition, where the startling auditory cue resulted in significantly lower reaction times during the transition from inspiration to expiration for the 1DI and FDS muscles. A Mann-Whitney U test was performed to compare reaction times between the different respiratory phases. Significant differences (p<0.05) between respiratory phases are indicated by connecting lines.

In the VRT condition (Figures 3A–3C), the sinusoidal fit revealed minimal modulation. For the 1DI muscle, the modulation was weak, with an amplitude of only 3.16 ms. FDS muscle showed a modulation amplitude that was slightly higher at 4.45 ms. Similarly, the biceps muscle displayed minimal modulation amplitude of 3.49 ms. These findings indicate weak not significant, respiratory phase-related modulation in the VRT condition.

In the VART condition (Figures 3D–3F), where a quiet auditory cue was added, sinusoidal fits indicated slightly stronger modulation. For the 1DI muscle, the fit showed a peak near the start of inspiration (I1), with a modest amplitude of 6.40 ms. The FDS muscle showed a peak just before the start of inspiration (I1), but the modulation was minimal, with an amplitude of -1.69 ms. The biceps muscle displayed a peak between the start of inspiration (I1) and the transition from the end of inspiration to the start of expiration (I3–E1), with an amplitude of 2.19 ms. Although no significant differences were observed between respiratory phases, the fit suggested subtle modulation across the respiratory cycle.

In the VSRT condition (Figures 3G–3I), the startling auditory stimulus elicited the strongest RT modulation. For the 1DI muscle, the sinusoidal fit showed that the RT trough occurred around the transition from the end of inspiration to the start of expiration (I3 to E1), with a significant modulation amplitude of 14.42 ms. There was a significant difference between RTs at the end of inspiration (I3) and the start of inspiration (I1) (p=0.025) and between the end of inspiration (I3) and the middle of inspiration (I2) (p=0.023). The FDS muscle showed modulation around the transition from the end of inspiration to the start of expiration (I3 to E1) with an amplitude of 9.27 ms. Lastly, the biceps muscle displayed modulation occurring around the same transition (I3 to E1) and an amplitude of 8.63 ms. There was a significant difference between RTs at the start of inspiration (I1) and the end of inspiration (I3) (p=0.005), the start of inspiration (I1) and the start of expiration (E1) (p=0.007), and between the start of inspiration (I1) and the end of expiration (E3) (p=0.022).

Figure 4 illustrates the StartReact effect, calculated as the difference in mean reaction times between the VART and VRST conditions. Statistical significance of peak and trough values was assessed using a bootstrapping method. For the 1DI muscle, the peak StartReact effect occurred at phase I3 (end of inspiration) with a value of 59.68, though this was not statistically significant (p = 0.079). Significant troughs were observed at phase E2 (middle of expiration), with a value of 32.48 (p = 0.028), while the trough at phase I2 (middle of inspiration) was not statistically significant (35.94, p = 0.094). These findings suggest a decrease in the StartReact effect during these respiratory phases, potentially indicating modulation of motor responses by respiratory rhythms.

**Figure 4:**
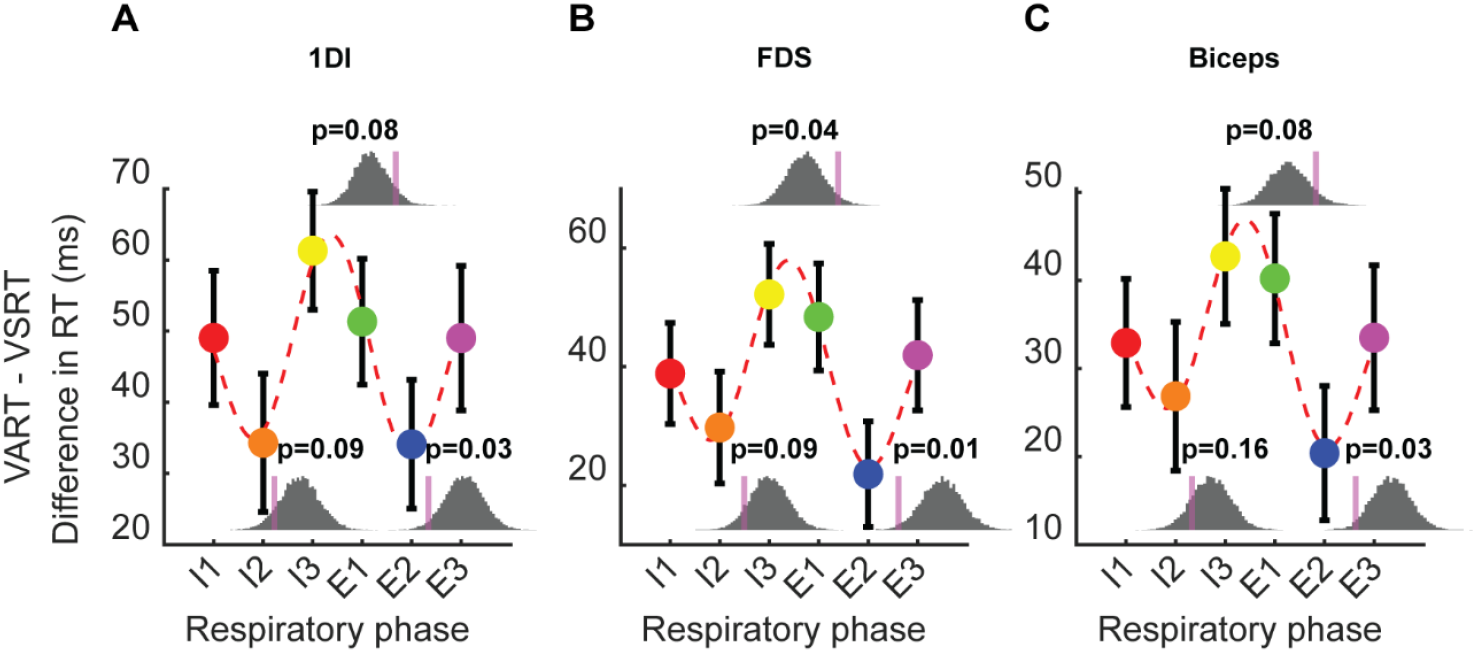
StartReact effect. This is represented for the three muscles: **(A)** first dorsal interosseous (1DI), **(B)** flexor digitorum superficialis (FDS), and **(C)** biceps. The mean and standard error of the mean (SEM) across the six respiratory phases (in blue inspiration and in red expiration) are shown along with the sinusoidal fit. Insets near respiratory phases I3-E1 (peaks) and phases I2 and E2 (troughs) illustrate the distribution of simulated peaks and troughs. The actual peak or trough value is marked as a vertical magenta line with its respective p-value.

For the FDS muscle, the peak value at phase I3 (end of inspiration) reached 53.39, achieving marginal statistical significance (p = 0.041). A significant trough was observed at phase E2 (middle of expiration), with a value of 23.08 (p = 0.015), while the trough at phase I2 (middle of inspiration) did not reach significance (28.56, p = 0.092). This pattern suggests a reduction in the StartReact effect during these phases, supporting the hypothesis of respiratory phase-specific modulation of motor pathways.

For the biceps muscle, the peak StartReact effect at phase I3 (end of inspiration) was 43.49, though not statistically significant (p = 0.084). A significant trough was found at phase E2 (middle of expiration), with a value of 21.15 (p = 0.027), whereas the trough at phase I2 (middle of inspiration) was not significant (26.10, p = 0.160). These results align with observations in other muscles, further suggesting that motor excitability is modulated by respiratory phase dynamics.

## Discussion

This study demonstrates that respiratory rhythms dynamically influence RST excitability, as evidenced by changes in RTs during the StartReact paradigm. We found that RTs were shortest during transitions from inspiration to expiration, particularly in the VSRT condition, where RST activation is maximized. These findings reveal a direct link between respiratory phase transitions and RST facilitation.

Our results, shown in Figure 2, align with previous studies (Honeycutt et al., 2013; Nonnekes et al., 2014; Dean and Baker, 2016; Singh et al., 2018; van Lith et al., 2018; Choudhury et al., 2019; Sangari and Perez, 2019; Smith et al., 2019; Sangari and Perez, 2020; Maslovat et al., 2021; Škarabot et al., 2022; Tapia et al., 2022; Maslovat et al., 2023; Walker et al., 2024), demonstrating a reduction in RTs across the conditions. The longest RTs were observed in response to unisensory visual stimuli (VRT). These RTs decreased when a quiet sound was presented alongside the visual stimuli (VART) and were further reduced when a startling sound accompanied the visual stimuli (VSRT). In this study, participants performed a multi-joint coordinated movement involving elbow flexion (biceps), wrist flexion (FDS), and a power grip (1DI). The RT reductions observed from VART compared to VRT, and from VSRT compared to both VART and VRT, were consistent across all three muscles.

The RTs observed in the VART condition compared to the VRT condition align with the well-documented redundant signals effect (RSE), wherein bisensory stimuli elicit faster responses than their unisensory components (Hershenson, 1962; Schröger and Widmann, 1998; Nidiffer et al., 2016). Traditionally, this speeding effect has been attributed to co-activation mechanisms, where neural integration of unisensory inputs enhances sensory-motor activation (Miller, 1982; Molholm and Foxe, 2010). However, recent findings by Shaw et al.(Shaw et al., 2020) challenge this interpretation, suggesting that much of the RSE observed in mixed-design paradigms may result from modality-switching and mixing costs. Specifically, frequent alternation between unisensory (e.g., visual) and bisensory (e.g., audio-visual) trials can slow RTs for unisensory conditions like VRT, thereby amplifying the perceived facilitation in bisensory conditions like VART (Wylie et al., 2009).

The consistent RT reductions in the startling condition VSRT compared to VRT and VART, across the biceps, FDS, and 1DI muscles support the idea that RST acts as a common pathway for distributing motor commands to both proximal and distal muscles (Peterson et al., 1979; Riddle et al., 2009; Zaaimi et al., 2018a). This mechanism enables rapid, coordinated multi-muscle responses, consistent with prior evidence of startle-evoked reliance on subcortical circuits (Valls-Solé et al., 1999; Carlsen et al., 2004; Nonnekes et al., 2014; Smith et al., 2019).

While corticospinal contributions to the observed RT reductions cannot be fully excluded (Alibiglou and MacKinnon, 2012; Marinovic and Tresilian, 2016), the significant RT differences between the VSRT and VART/VRT conditions strongly suggest a dominant role of the RST. This aligns with findings from Tapia and Baker (Tapia et al., 2022), who demonstrated that loud auditory stimuli suppress corticospinal activity but enhance reticulospinal activity in the early post-cue period, reinforcing the notion that the RST provides the majority of the descending drive during StartReact responses. Their computational modeling further supports the conclusion that a reticulospinal contribution of at least 60% is required to replicate the observed StartReact RT reductions.

To further isolate the RST’s contribution, we calculated the StartReact effect, focusing on the RT difference between the VART and the VSRT conditions to exclude general facilitation effects of auditory stimulation. This subtraction-based method provides a direct and interpretable measure of startle-evoked enhancement in motor output. The StartReact effect is significantly different from zero across all three muscles, confirming active recruitment by the reticulospinal system. Contrary to prior studies suggesting greater RST activation for proximal muscles (Maslovat et al., 2023), the gain (Fig 2C) was significantly higher for the intrinsic hand muscle (1DI) compared to the more proximal muscles (FDS and biceps). This could be due to a targeted increase in the RST drive towards the hand, optimizing its contribution to executing a power grip. Interestingly Perez and Baker found in spinal cord injury patients that RST input depended on the task with higher input during power grips than during precision grips (Baker and Perez, 2017).

The respiratory phase significantly influenced RTs, particularly in the VSRT condition, where RTs were shortest during the transition from inspiration to expiration (Phases 3–4) for the 1DI and FDS muscles. This IE transition phase (von Euler and Trippenbach, 1975; Hülsmann et al., 2021) as well as the reverse (EI) represents critical moments in the body’s physiological rhythm, with notable effects on cortical and motor excitability (Zelano et al., 2016; Nakamura et al., 2018). Similarly, these respiratory transitions seem to modulate RST. As shown in Figure 4, it is evident that RST gain increases during transitions between inspiration and expiration, as well as near the transitions from expiration to inspiration and significant decreases in RST gain are observed mid-expiration across all three muscles. This pattern highlights the dynamic modulation of RST motor pathway excitability by respiratory phase.

By pairing VNS with task-specific rehabilitation, studies have shown significant improvements in motor function, suggesting that VNS triggers long-term plasticity in motor circuits (Hays et al., 2016; Engineer et al., 2019; Dawson et al., 2021). While cortical reorganization is frequently proposed as a primary mechanism of VNS-induced recovery (Khodaparast et al., 2014), a more anatomically proximal motor circuit may also play a role. Specifically, the reticular formation and its motor output, the RST, may be modulated by VNS. Vagus nerve afferents project to the nucleus tractus solitarius in the brainstem, which connects to the reticular formation (Peyron et al., 1996; Ruggiero et al., 2000). By activating the reticular formation, VNS could influence RST activity, potentially driving motor recovery. Given the vagus nerve’s role in regulating breathing rhythms and the transition between inspiration and expiration (von Euler and Trippenbach, 1975; von Euler, 1983), One hypothesis is that VNS could harness the neuromodulatory effects of respiration to enhance RST activity and strengthen its contribution to motor recovery. While VNS’s direct effects on the RST remain to be demonstrated, this pathway represents a promising target for leveraging respiratory-driven neural plasticity to optimize motor recovery.

The recent study by Choudhury et al. (Choudhury et al., 2020) presents an intriguing application of RST modulation in motor rehabilitation. Their wearable device uses auditory-motor pairing to induce plasticity in the reticulospinal system improving upper-limb function in chronic stroke patients. Our findings complement this approach by highlighting the importance of respiratory-phase transitions in modulating RST excitability. Specifically, targeting stimulation to coincide with the transition from inspiration to expiration could optimize the timing of therapeutic interventions, potentially enhancing motor recovery outcomes.

In summary, this study demonstrates that respiratory rhythms significantly influence reticulospinal tract excitability, with transitions between respiratory phases serving as critical moments for modulating motor pathway activity. By combining these insights with existing evidence of RST plasticity and its role in motor recovery, our findings provide a foundation for future research aimed at integrating respiratory-phase-dependent modulation into neurorehabilitation strategies. Targeted interventions, such as respiratory-phase-aligned stimulation using wearable devices leveraging auditory-motor pairing, may harness these mechanisms to enhance motor recovery following corticospinal lesions. These advances could pave the way for personalized therapies that align with the body’s natural rhythms to maximize functional outcomes.

## Acknowledgement

We would like to extend our sincere thanks to all the participants for their time and valuable contributions to this study. Your contributions were crucial to the success of this project.

